# The controlled spectrum of plant form and function

**DOI:** 10.64898/2026.07.27.740885

**Authors:** Milad Rahimi-Majd, Alistair Leverett, Johannes Kromdijk, Zoran Nikoloski

## Abstract

Dimension reduction techniques with plant traits have suggested the existence of independent acquisitive-conservative spectra in leaves and roots across diverse plants. However, this finding may be due to third-variable effects that can alter the fundamental dimensionality of plant trait variation. Using a dimension reduction approach that controls for third-variable effects, we show that the acquisitive-conservative axis is not confined within an organ, but acts at the whole plant level.

## 1 Introduction

Vascular plants are fundamental drivers of Earth’s ecosystems, shaping the flow of energy and matter while affecting the balance between oxygen and carbon dioxide [1]. Rapid global change places unprecedented pressure on biodiversity and ecosystem functioning, underscoring the need to understand how plants respond to complex, dynamic environments. The remarkable structural, morphological, and functional diversity is reflected in a wide array of plant traits associated with ecological strategies and ecosystem processes [2]. To better understand how plants interact with ecosystems, efforts have focused on identifying the fundamental dimensions of plant trait variation [1, 2, 3, 4, 5, 6, 7]. Dimension reduction techniques have pinpointed key dichotomies and trade-offs among groups of traits, enabling us to model plant plasticity and forecast ecosystem responses for future climate scenarios [4, 5, 6, 7].

The world-wide leaf economics spectrum (LES) offered an early hypothesis on the dimensionality of plant traits. The theory of the LES posits that a single acquisitive–conservative axis from a principal component analysis (PCA) captures the majority of trait variance. Based on the rate of return on carbon, nutrient, and water investments, this axis is interpreted as a fundamental trade-off between carbon investment (growth) and leaf lifespan [8, 9, 10]. This framework may extend beyond leaf level traits, and a plant economic spectrum has been hypothesised, in which roots, stems, and leaves are coordinated in their rates of resource acquisition [11].

However, multiple dimensions of variation have been uncovered when traits from other organs are jointly considered. For example, the global spectrum of plant form and function (GSPFF) [1] identified two orthogonal dimensions in the above-ground trait space. The first axis captures the dichotomy between woody and herbaceous species, based on the plant size, whereas the second axis represents the LES. Furthermore, analysis of root traits identified two dimensions across a root economics space (RES) [4]. The first axis represents an acquisition-conservation gradient, conceptually analogous to the LES. The second axis captures a collaboration gradient, defined by a trade-off between direct soil exploration, through increased specific root length, and reliance on mycorrhizal partners, via greater carbon investment and thicker root diameters.

Whilst the GSPFF and RES exhibit multidimensionality, both contain an acquisitive-conservative axis. Alignment of these axes when the above- and below-ground traits are jointly analyzed would provide evidence in favor of a plant economic spectrum for nutrient acquisition, that has remained elusive. For example, the Unified Plant Functional Space (UPFS) [6] found that four dimensions were needed to explain 76% of trait variance: two representing the LES and plant size and the other two representing the RES. This finding suggests that nutrient acquisitive-conservative spectra in leaves and roots are independent. A recent analysis based on newly acquired data found orthogonality between leaf and root nitrogen, further supporting the hypothesis that these traits are independent [12]. Despite criticism of the UPFS [5, 13], this framework has remained the most convincing explanation for plant form diversity and appears to be robust to data augmentation [7].

## 2 Results

### 2.1 Third-variable effects are ubiquitous in triads of plant traits

Whilst the aforementioned analyses suggest orthogonality between leaf and root nitrogen, these approaches are based on PCA, which does not inherently consider the influence of third-variable effects on pairwise trait correlations. Specifically, the presence of such third-variable effects among traits is expected to shape the core patterns of trait covariation and consequently, the dimensionality inferred by PCA. To address this issue, we sought to account for third-variable effects before conducting PCA, to reassess both the dimensionality of data and the degree to which acquisitive-conservative axes in the leaves and roots are aligned when third-variable effects are explicitly considered. To relate our work to previous research, we used the data set from Ref. [6], which substantially overlaps with those employed in the studies discussed above. This data set includes ten above- and below-ground plant traits, namely: specific stem density (*ssd*), leaf area (*la*), leaf nitrogen concentration (*ln*), leaf mass per area (*lma* = 1*/sla*), seed diaspore mass (*sm*), plant height (*ph*), root diameter (*D*), root tissue density (*RTD*), root nitrogen concentration (*N*), and specific root length (*SRL*) (see Methods).

We first examined the extent to which the direct Pearson correlation (*r*) between each pair of traits deviates from their corresponding partial correlation (*r*_*c*_) values when controlling for one co-variate (see Methods). We denote the absolute difference between the two correlation coefficients, while considering their significance, for each pair of traits, as Δ = |*r* − *r*_*c*_|. We observed general deviations between *r* and *r*_*c*_ for different pairs of traits when controlling for another trait as a co-variate (Fig. 1a). In 38 cases, the deviations (Δ) reached values larger than 0.1, suggesting that *r* is strongly affected by other traits (Fig. 1b).

**Figure 1.**
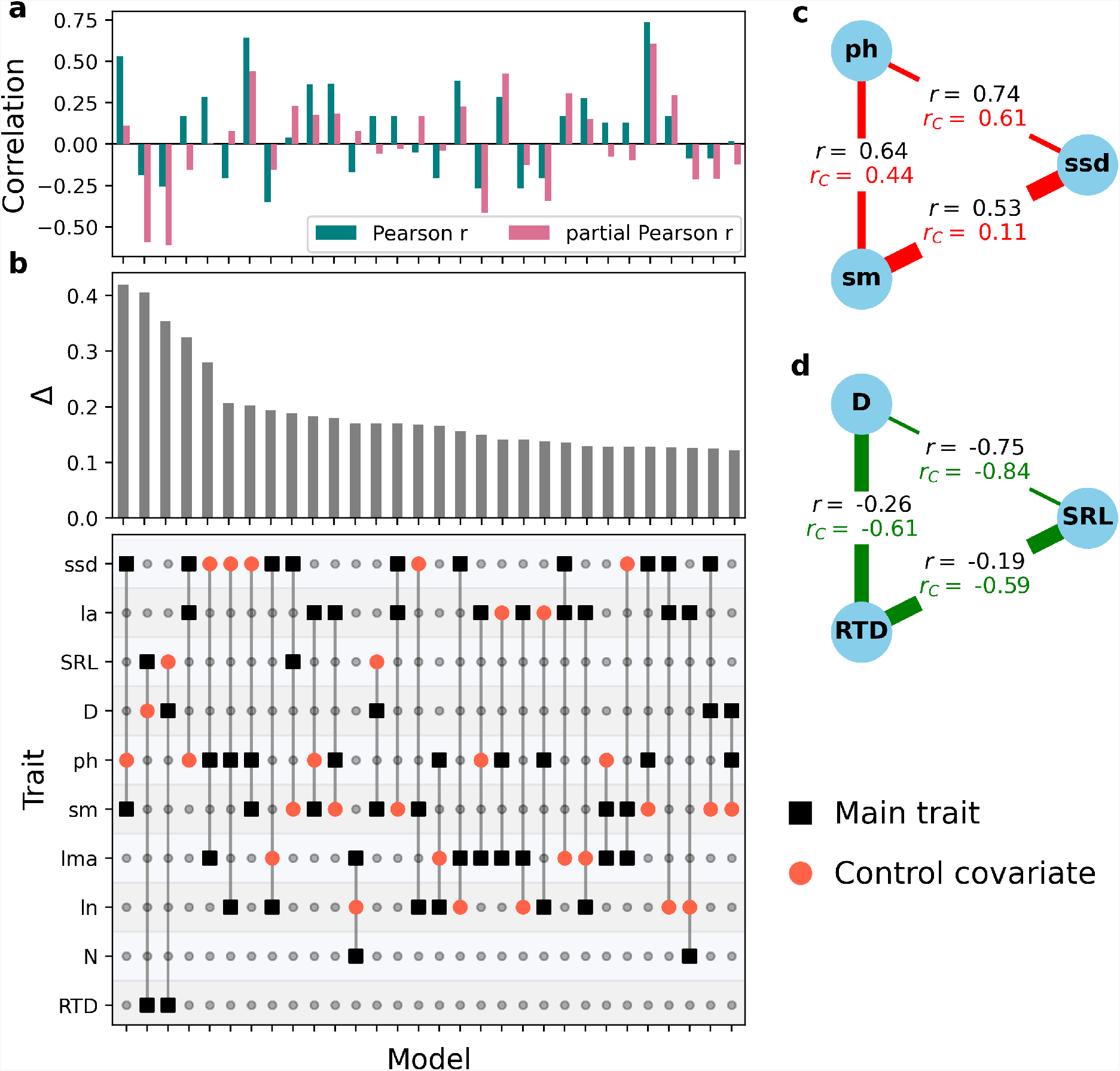
Pearson correlations between pairs of plant functional traits are altered upon control for a third trait. **a**, Pearson (*r*) and partial (*r*_*C*_) correlation coefficients for pairs of plant traits, computed for each model defined by the trait–covariate matrix, shown below. Each column in the matrix represents one model, with black squares denoting the two focal traits whose correlation is investigated with and without inclusion of a control covariate, marked with orange circle. The analyzed ten traits comprise: specific stem density (*ssd*), leaf area (*la*), leaf nitrogen concentration (*ln*), leaf mass per area (*lma*), seed diaspore mass (*sm*), plant height (*ph*), root diameter (*D*), root tissue density (*RTD*), root nitrogen concentration (*N*), and specific root length (*SRL*). **b**, Absolute difference (Δ = |*r* − *r*_*C*_|) between direct and partial Pearson correlations (with significance constraints described in Methods), reflecting the extent to which controlling for a covariate alters the Pearson correlation between two traits. The plot includes the 30 models with the largest Δ values. **c**-**d**, Example partial Pearson correlation networks for selected triplets of traits, illustrating how partial correlations can strengthen (green edges) or weaken (red edges) Pearson correlations by controlling for the third trait in the triplet. Edge width scales with the magnitude of Δ. the three-dimensional latent space, we defined a composite direction by averaging the loadings of the corresponding traits. The resulting three dimensions of trait variation exhibited near-perfect orthogonality, with pairwise angles ranging from *θ* = 88.1^*°*^ to 92.8^*°*^ (Fig. 2c,e).

This observation was particularly evident for a triplet of above-ground traits associated with plant size, i.e., *ph, sm*, and *ssd* (Fig. 1c), as well as for a triplet of below-ground traits, i.e., *D, SRL*, and *RTD* (Fig. 1d). We observed that the magnitude of the weak correlation between *SRL* and *RTD* increased markedly when controlling for *D* (Δ = 0.40), and a similar pattern was found for the correlation between *D* and *RTD* controlled by *SRL*. However, controlling for *RTD* had only a minor effect on the strong negative correlation between *D* and *SRL* (Fig. 1d). Conversely, controlling the correlations for pairs of traits in the triplet *ph, sm*, and *ssd* led to a decrease in the magnitudes of their correlations; for instance, the correlation between *sm* and *ssd* strongly decreased when controlling for *ph* (Δ = 0.42) (Fig. 1c). These observations indicate that the covariation does not arise from a direct dependence, but rather from underlying relationships with a common variable [14].

### 2.2 Specific combinations of plant traits capture the majority of third-variable effects

Having established the presence of third-variable effects, we then sought to identify a minimal subset of the analyzed traits that maximizes the total deviations (denoted by *S*_Δ_) between direct and partial correlations across all possible combinations of pairs and extension with a control variable (see Methods). The result of this analysis will provide candidates for third-variable effects that need to be controlled. This analysis revealed that a subset consisting of two above-ground traits, *ssd* and *la*, together with either *D* or *SRL* as below-ground traits, accounted for 92% and 91% of the total *S*_Δ_, respectively. We note that *ssd* and *la* are traits that do not align well with the two major dimensions in the GSPFF (see Ref. [1]). In fact, these traits are correlated with both plant size and the LES, thereby affecting the majority of the traits. In addition, the selected variables recapitulate the inverse relationship of *D* and *SRL* with *RTD* that was theoretically established using the cylindrical geometry of roots [15], however, without the need for additional assumptions about the independence of *RTD*.

### 2.3 Conditional PCA of lower dimensionality points at cross-organ integration

To reveal the controlled spectrum of plant form and function, next we performed a conditional PCA. This approach involved regressing each trait against the three control traits (i.e., *ssd, la*, and either *D* or *SRL*) and performing a PCA with the resulting residuals (see Methods). The outcome of this approach, for both scenarios, was a PCA space in which 68% of variance was captured by three major axes (Fig. 2a-c, e). The first dimension of the leaf traits captures the LES, and is defined by the opposing loadings of *ln* and *lma* in the PC1–PC2 plane (Fig. 2a,c,e). We also found that *N* is strongly coordinated with *ln* within the same plane. The second dimension represents the RES, defined by the loadings of *D* (or *SRL*) and *RTD* (Fig. 2a,c,e), together with the well-known trade-off between *D* and *SRL* (Fig. 2f) [4]. This finding suggests an extended trade-off dimension for RES, reflecting a dichotomy between plants with conservative roots, characterized by higher nutrient investment through increased *RTD*, and plants with acquisitive roots, which follow one of two alternative strategies: (i) outsourcing nutrient acquisition via mycorrhizal partners through increased *D* or (ii) a “do-it-yourself” strategy via increased *SRL* (Fig. 2c,e-f) [4]. Finally, the third dimension captures the plant size variation described by the variables *ph* and *sm* (Fig. 2b,c,e). Importantly, this axis of variation preserves the dichotomy between woody and herbaceous species (Fig. 2d) in accordance with the previously described plant size gradient [1]. This highlights the ecological relevance of the controlled plant size dimension despite the exclusion of *ssd* and *la* from the main variables. For each major axis of variation (i.e., LES, RES, and plant size), in

**Figure 2.**
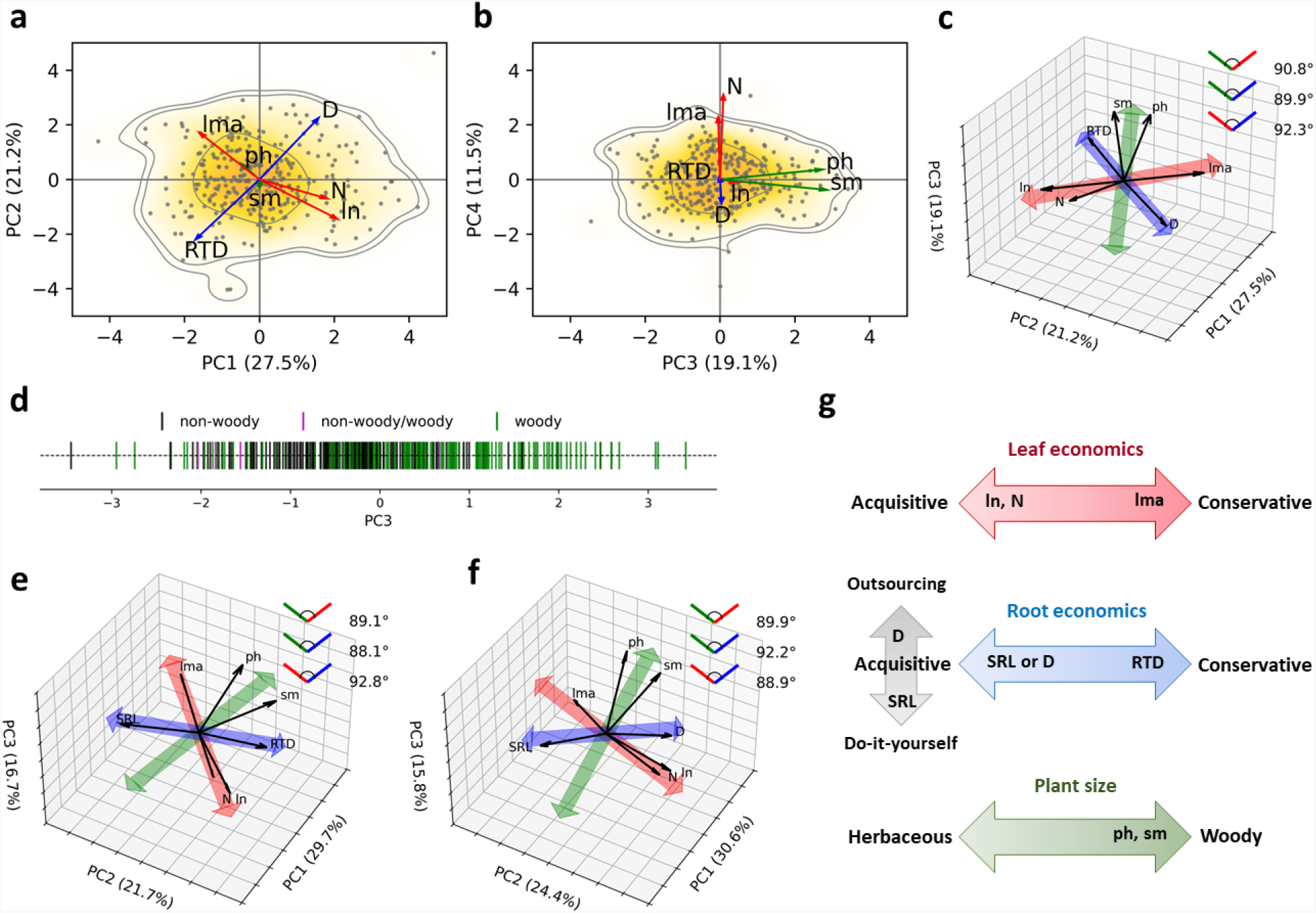
**a**-**b**, Projection of plant species in the functional trait space defined by conditional principal component analysis (PCA) axes, showing the first four principal components. The analysis was performed on the residuals of seven traits—leaf nitrogen concentration (*ln*), leaf mass per area (*lma*), seed mass (*sm*), plant height (*ph*), root diameter (*D*), root tissue density (*RTD*), and root nitrogen concentration (*N*)—after statistically controlling for three traits: specific stem density (*ssd*), leaf area (*la*), and specific root length (*SRL*). Dots represent species, and the color gradient (white–yellow–orange) indicates occurrence probability, from the lowest to the highest, with contour lines delineating the 0.5, 0.95, and 0.97 quantiles. Coloured arrows indicate the direction and weighting of scaled trait loadings (red: leaf economics, blue: root economics, and green: plant size), illustrating the main dimensions of covariation among the analyzed traits. **c**, Three-dimensional biplot representation of the first three principal components (PC1–PC3) based on scaled trait loadings. Black arrows denote the original trait loadings, labeled by trait name. Colored double-headed arrows indicate the major composite directions of three trait groups, obtained as the mean direction of the loadings within each group: red (*lma, ln, N*), blue (*RTD, D*), and green (*ph, sm*). The legend displays the pairwise angles (in degrees) between these community-level directions. **d**, Scatter plot of species along PC3, with vertical bars colored by woodiness category, showing a general separation between woody and herbaceous species. **e**, As in panel **c**, with *D* instead of *SRL* treated as a control trait. **f**, As in panel **e**, with *D* instead of *RTD* in the main group of traits, and *RTD* excluded from the analysis. **g**, Conceptual representation of the controlled spectrum of plant form and function.

## 3 Discussion

In summary, using an approach based on conditional PCA with data on combined plant above- and below-ground traits, we found that the resulting controlled trait space exhibits lower dimensionality than existing models [6, 7] that use the same traits. This controlled trait space reveals three principal dimensions of variation: (i) an above-ground size gradient, (ii) an extended RES, capturing the gradient from conservative to acquisitive roots, the latter employing either ‘do-it-yourself’ or ‘outsourcing’ strategies, and (iii) an extended LES, due to alignment of *N* with the direction of *lma*, and *ln* (Fig. 2g). Taken together, these analyses suggest that a component of the nutrient acquisitive-conservative axis of variation may not be confined to specific organs but rather may act at the level of the whole plant.

## 4 Materials and Methods

### 4.1 Data set

To investigate the relationships among plant functional traits, we utilized the vascular plant data set from Carmona et al. (2021) [6], available at https://doi.org/10.6084/m9.figshare.13140146. This data set substantially overlaps with those employed in several highlighted studies in this field, including the references [1, 4, 5, 7, 13, 16]. This data set includes measurements for six above-ground functional traits derived from the TRY Plant Trait Database [17]: leaf area (*la*), specific leaf area (*sla*), leaf nitrogen concentration (*ln*), specific stem density (*ssd*), seed (diaspore) mass (*sm*), and plant height (*ph*). In all our analyses, we consider the inverse of the *sla* as leaf mass per area, *lma* (i.e. *lma* = 1*/sla*). Four below-ground traits were also analyzed, extracted and integrated from the GRooT global database [18]: root diameter (*D*), specific root length (*SRL*), root tissue density (*RTD*), and root nitrogen concentration (*N*). In total, the dataset comprises approximately 40,000 species, among which 2630, 748, and 301 species have complete information for above-ground, below-ground, and all ten traits, respectively. All analyses were conducted on the subset of 301 species with complete data for all ten traits. The above-ground traits were log-transformed and standardized using z-transformation, while the below-ground traits had already undergone log-transformation in the original, source data set.

### 4.2 Partial correlation analyses

Partial correlation (*r*_*c*_) measures the direction and magnitude of a relationship between two random variables *X* and *Y* , after eliminating the effect of one or more variables (i.e., covariates). Considering the covariate variable as *Z*, we used the method defining the partial correlation between two variables *X* and *Y* conditioned on *Z* as:

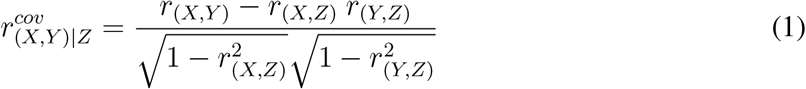

where *r* and *r*^*cov*^ indicate the direct and partial correlations between the given variables, respectively [19]. In this work, all the partial correlation analyses were calculated using the python package *Pinguan* [20].

### 4.3 The minimal set of covariate traits

Having 10 traits, both Pearson (*r*) and partial (*r*_*c*_) correlations for the main variables of all the 360 possible triplets of traits (two interchangeable main variables and a control covariate) were assessed. From these combinations, in 189 cases, both correlation coefficients (*r* and *r*_*c*_) were statistically significant at the *p <* 0.05 level. In 126 cases, neither correlation reached statistical significance, while in the remaining cases, only one of the two correlations was found to be significant.

By excluding the cases in which both correlations were non-significant from the analysis, we quantified the deviation between the two correlations using their absolute difference as:

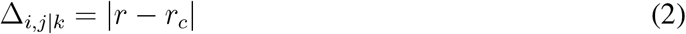

where *i* and *j* denote the main pair of traits, and *k* represents the control covariate. To avoid spuriously large Δ values in cases where only one of the correlations was significant, the non-significant correlation was set to zero if its sign was opposite to that of the significant correlation.

We defined the weighted total difference between the two correlations across all analyzed triplet combinations of traits as:

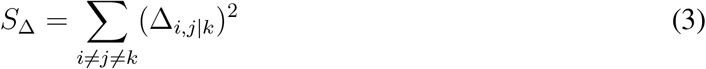

where *i* and *j* span all possible pairs of traits for which at least one of the corresponding correlations (*r* or *r*_*c*_) is significant, and *k* denotes each of the remaining traits considered as the covariate variable. Squaring the Δ values ensures that larger control effects contribute more strongly to the overall value of *S*_Δ_.

### 4.4 Conditional principal component analysis

Conditional Principal Component Analysis (PCA) was performed to examine the latent structure of the data while accounting for potential spurious effects of specific control variables. In this procedure, each of seven main variables (i.e., four above-ground and three below-ground traits) was first linearly regressed on the set of three control variables (i.e., two above-ground and one below-ground trait) using an Ordinary Least Squares (OLS) model. The corresponding residuals were extracted to represent the portion of variance having eliminated the linear part explained by the control variables. The resulting residual matrix was standardized (zero mean, unit variance) and subjected to principal component analysis (PCA). Importantly, this normalization process ensures that the analysis captures the relative trade-offs and couplings between traits, rather than their controlled variance magnitudes. The OLS model implementation and PCA analysis were performed using Python package *Scikit-learn*, version 1.6.1 [21].

### 4.5 Definition of loading dimensions

Composite dimensions were defined to capture the dominant directional trends shared among related functional traits in the PCA space. For each composite dimension, the loading vectors of the constituent features (taken from the first three principal components) were converted to unit vectors and averaged within three predefined groups: {−*LMA, ln, N*}, {*ssd, ph, sm*}, and {*RTD*, −*D*, −*SRL*}. The negative signs indicate that the corresponding loading vectors were flipped in orientation prior to averaging, ensuring that all vectors within a group were directionally aligned. The angular separation between any two composite vectors, **v**_1_ and **v**_2_, was computed by normalizing each to unit length and evaluating

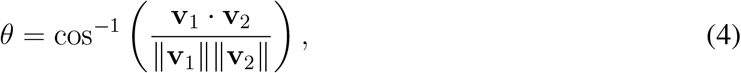

where *θ* is expressed in degrees.

### 4.6 Occurrence probability

To quantify and visualize the occurrence probability distribution of sample scores in the principal component (PC) planes, a non-parametric Kernel Density Estimation (KDE) method was employed using a Gaussian kernel. The KDE was performed in two dimensions, corresponding to each selected pair of principal components, to estimate a continuous probability density distribution of sample occurrence across the PC plane. The estimated density was normalized such that the total cumulative probability integrated to 1. To delineate the most probable regions of sample occurrence, contour lines were computed for cumulative probability quantiles of 0.50, 0.95, and 0.97, corresponding to the 50%, 95%, and 97% confidence regions, respectively. The KDE computation was performed using the *SciPy* Python library [22], version 1.15.2.

## 5 Data availability

The code supporting our analyses are available at github.com/MRahimiMajd/Plant controlled spectrum. The data set used in this study is available using the link: https://doi.org/10.6084/m9.figshare.13140146 [6].

## 7 Author contributions

M.R.-M. conceptualized the study, performed the computations, analyzed the data, interpreted the results, and wrote the paper. A.L. analyzed the data, interpreted results, and wrote the paper. J.K. interpreted results and and wrote the paper. Z.N. conceptualized the study, analyzed the data, interpreted results, wrote the paper. All authors contributed to finalizing the manuscript.

